# ColabPCR: A validated Google Colaboratory Notebook for Reproducible and Precise Primer Design

**DOI:** 10.64898/2025.12.23.696274

**Authors:** Mauricio J. Lozano, Dardo Dallachiesa, Gernot Beihammer, Jennifer Schwestka, Julia Koenig-Beihammer, Eduardo José Peña

**Affiliations:** IBBM, Instituto de Biotecnología y Biología Molecular, Fac. Cs. Exactas Universidad Nacional de La Plata, CONICET-CCT La Plata; Ergo Bioscience SAS, Av Belgrano 758, Sunchales, PC 2322, Santa Fe, Argentina; NOVABIOMA Biomanufacturing FlexCo, Vienna, Austria; Austrian Centre of Industrial Biotechnology (ACIB), Graz, Austria

**Author notes:** Correspondence: Mauricio J. Lozano,; Eduardo José Peña. Authors contributed equally to this work (Mauricio J. Lozano, Dardo Dallachiesa).

**Keywords:** Gene expression, Polymerase chain reaction, Primer design, Virtual PCR, Primer3

## Abstract

The Polymerase Chain Reaction (PCR) method often has a lower success rate when amplifying specific genomic regions in eukaryotic genomes. This is frequently due to the non-specific annealing of primers at various genomic locations. To address this issue, we created ColabPCR, a program specifically designed to optimize the selection and evaluation of primers for a defined genomic region. ColabPCR refines primer length and melting temperature parameters through the utilization of Primer3 software, and it quantifies the number of potential off-target binding regions within the target genome via BLASTn analysis. Furthermore, the program facilitates the integration of restriction enzyme recognition sites at the 5’ termini of primers and provides a mechanism to confirm the absence of such sites within the intended amplification region. ColabPCR centralizes all these functionalities within a single interface, utilizing Google Colab’s computational resources to ensure high performance and accessibility without requiring local software installation. We rigorously validated ColabPCR by designing primers for promoter and terminator regions within the *Daucus carota* (DCARv2, DH1v3) reference genome. Our findings unequivocally demonstrate significantly enhanced success rates, particularly when primers exhibiting off-target binding are excluded from the primer design process. In summary, ColabPCR offers a user-friendly and powerful solution that simplifies and enhances the primer design and evaluation workflow, leading to increased accuracy and success in molecular biology experiments.

## Introduction

In molecular biology, effective primer design is crucial for successful polymerase chain reaction (PCR) amplification of specific DNA regions. Key requirements for primers include similar melting temperatures (Tm), balanced G/C content, and the prevention of self-complementarity and hair-pin structures ^1^. Various software tools are available for designing viable primer sets from a given target DNA sequence taking into account these critical factors. Primer3 ^2^ is a widely used software program for designing PCR primers, hybridization probes and sequencing primers. It takes a DNA sequence and design parameters as input and outputs primer sequences that meet the specified criteria, such as desired length, melting temperature, GC content, and product size. The program uses advanced thermodynamic models for melting temperature calculations and for estimating the propensity of primers to hybridize with other primers or to hybridize at unintended sites in the template.

Another requirement is that Primers must specifically amplify the target region. The design of specific primers typically involves two stages. First, primers that flank the regions of interest are generated, either manually or with specialized software. Primer design software requires the input of the target region, which is typically defined by the user and provided manually as a genomic DNA sequence fragment. Then, the specificity of these primers is evaluated by searching nucleotide sequence databases using tools like BLAST, to identify possible undesired targets ^1,3–5^. Off-target primer binding can occur even with several mismatches, reducing the yield of the PCR reaction due to nonspecific amplification and primer consumption by linear amplification of non-target sequences.

Several tools like Primer-Blast (QuantPrime ^6^, PRIMEGENS ^7^) design primers using Primer3 software and then select specific primers using megablast searches against a genomic database. These programs use different criteria to select specific primer pairs, but do not provide further information about the nonspecific binding that could be useful when non-specific primers are found. Other tools for primer design are oriented either to expand the target range for a single primer set ^8^ or to design microbe- and group-specific primers ^9^.

This work introduces ColabPCR, a novel primer design tool offering functionalities akin to Primer-Blast, yet incorporating specific features for designing primers for PCR amplicon cloning and evaluating target specificity. Given an NCBI assembly accession number, ColabPCR retrieves the corresponding genome, extracts the target sequence using a specified locus tag or genomic coordinates, and generates a restriction enzyme analysis report to support construct design. The platform further enables the incorporation of custom oligonucleotide sequences at the 5′ end of primers for downstream applications, designs primer sets for amplification of the selected region using Primer3, and assesses their specificity with BLASTn ^10^, reporting predicted off-target binding sites to guide selection of the optimal primer pair. We tested our tool by designing different sets of primers for the amplification of promoter and terminator regions of carrot (*Daucus carota*) obtaining an overall higher success amplification rate, compatible with automated platforms and high throughput cloning projects.

## Implementation

ColabPCR is a python notebook implemented on Google Colab that uses NCBI datasets ^11^ program to download the target genome in genbank format, Primer3 ^2^ for the design of primers in a specified genomic region, and lastly, the latest version of Blastn ^10^ to perform a virtual PCR. Sequence manipulation is carried on using Biopython ^12^ and table management using Pandas ^13^.

The first step is downloading the target genome (presets for *Daucus carota* genomes are included) by NCBI’s assembly^14^ accession number. Options for manually downloaded files are also available. Additionally, a custom sequence can be used as a target instead of a genomic region, by simply copying and pasting the sequence in the corresponding form field. Next, the region where primers are to be designed is extracted from the target genome. Here four modes can be selected: promoter, terminator, gene or region. All extraction modes require the specification of the genomic replicon or chromosome index. In the region mode, the DNA sequence defined by start, end positions is extracted from the specified chromosome or replicon. In the gene, promoter and terminator modes, the locus tag must be provided. For gene extraction the locus tag must correspond to the gene feature, while for the promoter and terminator modes the coding sequence (CDS type feature) locus tag is required. A flexibility option can be used to allow the variation of the start or end positions in a specified number of nucleotides. This option is for both sides unless the ends are set to fixed with the fixedNear or fixedFar options. The effect of the fixedNear and fixedFar options is different in the region, gene, promoter, and terminator modes. In the first three modes, the FixedNear option will fix the location of the primer in the start codon side (right side in the region mode) of the gene, while in terminator mode, the primer fixed will be the one located in the stop codon side of the gene. This location is relative, and depends on whether the gene is coded in the plus or minus strand.

Once the target region is extracted, a restriction enzyme (RE) analysis is conducted. This analysis first identifies restriction enzymes (REs) that cut within the target region. Subsequently, it offers the option to incorporate oligonucleotides, such as RE recognition sequences, at the 5’-end of the primers for downstream applications like PCR cloning. All the REs supported by Biopython are available.

Next, Primer3 is used to design 5 primer sets using the default options, and setting the 5’-added oligonucleotides, the product length range, and whether the primers should be fixed on either side of the target region (in such a case, only primer length would change). As a result, a table with the best 5 primer sets is generated.

The virtual PCR module employs BLASTn to identify primers within the target genome, allowing for mismatches and predicting potential PCR products. ColabPCR uses BLASTn with the following default options: -task blastn-short -word_size 4 -evalue 1 to search for short sequences and allow mismatches in the results. The BLAST results are analyzed and hits are classified by the number of mismatches, the orientation of forward and reverse primers and whether the primers are able to produce a PCR product of less than 20.000 nucleotides. This module can be operated with either pre-designed or custom primer sets.

## Results and discussion

### ColabPCR

Here we present ColabPCR, a python notebook designed to allow automatization and to improve the reproducibility of primer design for molecular biology experiments. The program can be run as a jupyter notebook or in google Colab (https://github.com/maurijlozano/ColabPRC). Figure 1 and Figure 2, show a schematic representation of the algorithm jointly with the Graphical User Interface (GUI) of ColabPCR with all the available options. The notebook works by running all the code blocks in the specified order. The first and second blocks install all the required dependencies –including BLAST, Datasets, and Biopython– and define the python functions for the target region extraction and virtual PCR blocks respectively.

**Figure 1.**
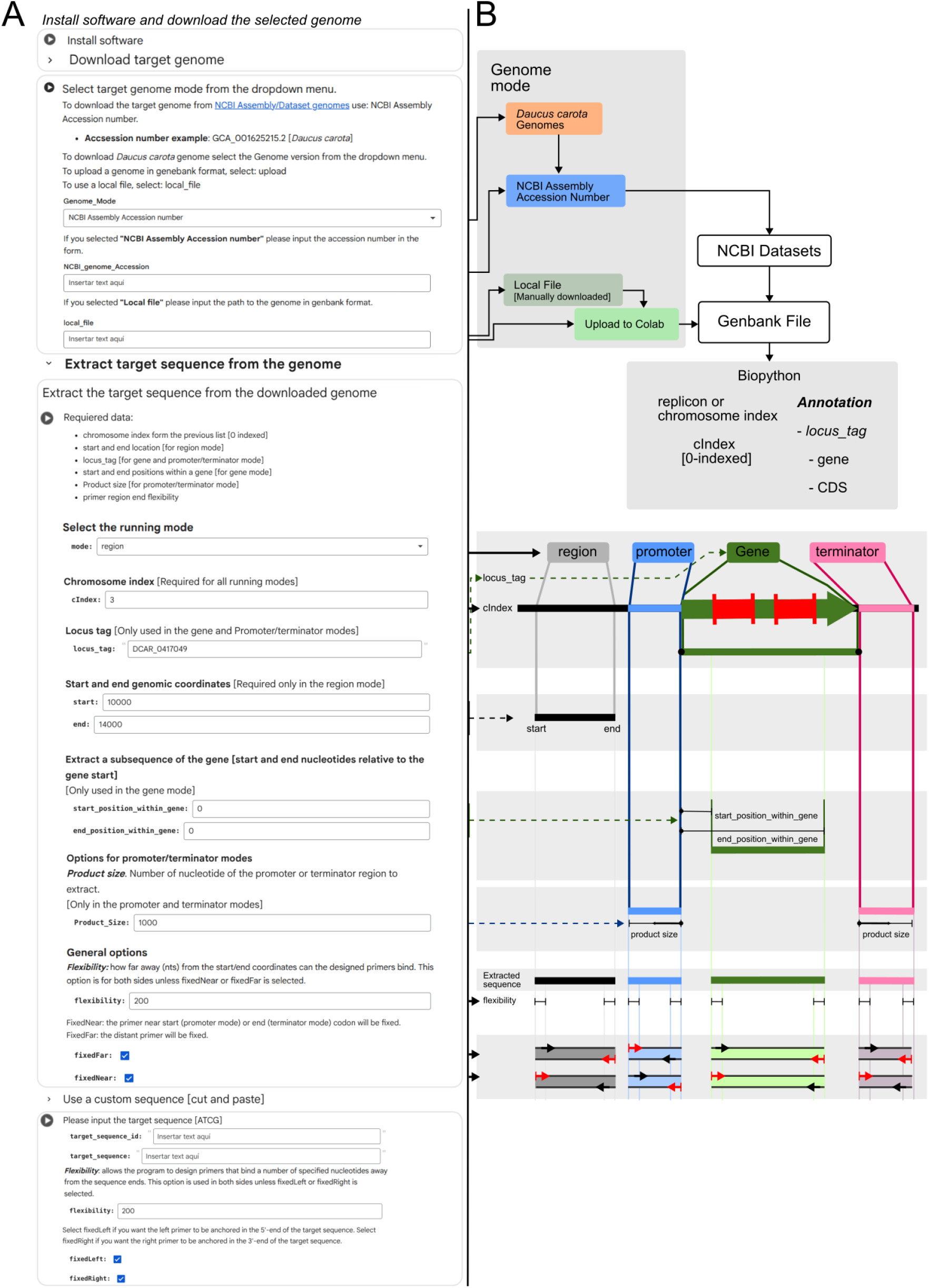
ColabPCR Options and algorithm. A. The form GUI provided by google Colab with all the available options. Dependencies installation, genome downloading and template region extraction. B. A scheme of the algorithm used by ColabPCR.

**Figure 2.**
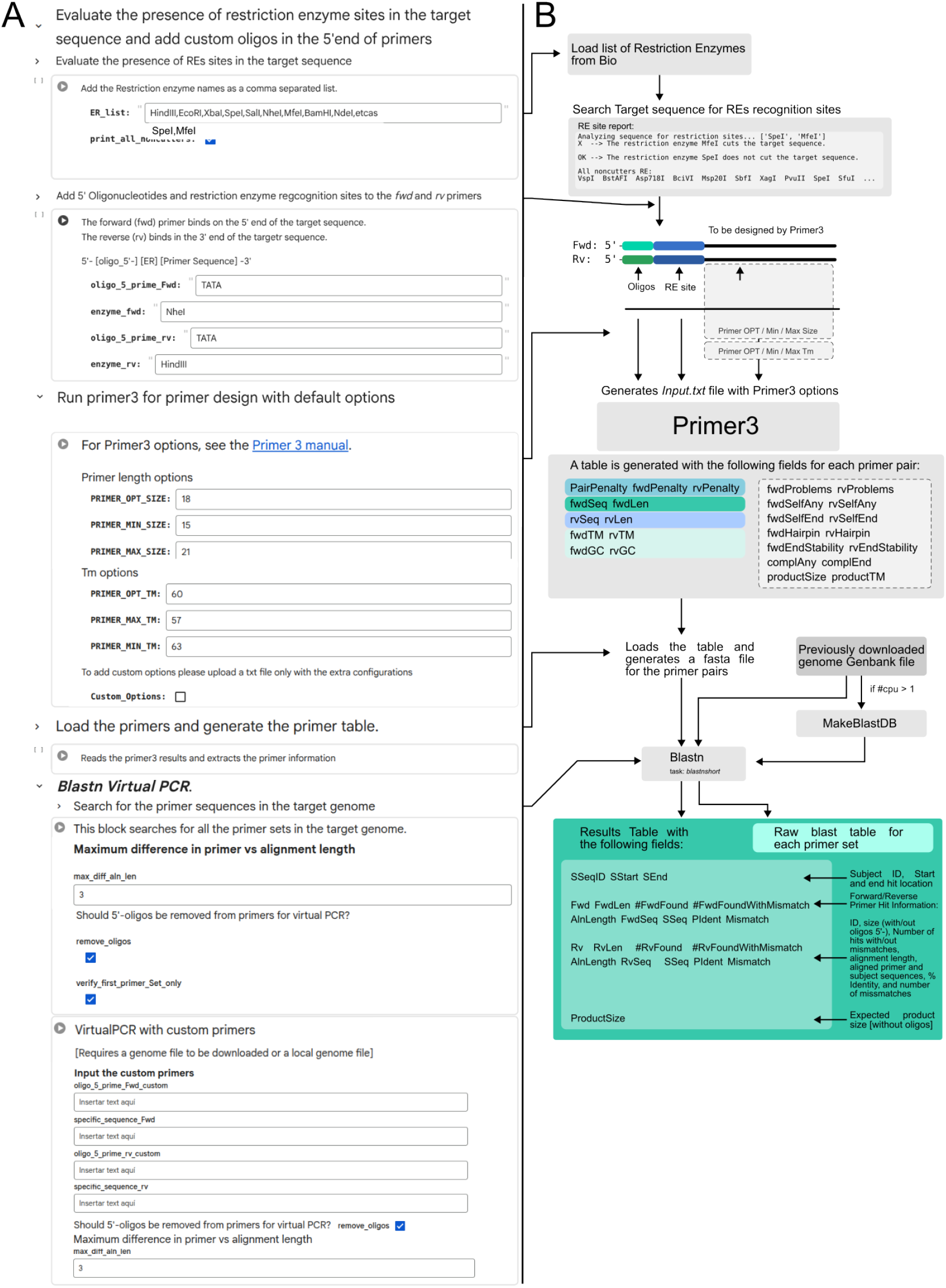
ColabPCR Options and algorithm. A. The form GUI provided by google Colab with all the available options. RE analysis, add custom 5’-oligonucleotides to primers and Virtual PCR. B. A scheme of the algorithm used by ColabPCR and the provided results.

Next, a genome mode must be selected. The available modes are *Daucus carota* genomes in two different versions, input a NCBI Assembly/Datasets accession number, and upload –or provide a path to an already uploaded– genbank file. Biopython is then used to load the genome, report the number of replicons/chromosomes found and their corresponding index, and extract a specified region using the information provided by the user. Four extraction modes are available. The *region* mode extracts a genomic DNA sequence using the chromosome index (cIndex, 0-indexed) and the start and end coordinates. The *gene* mode uses the *cIndex* and *locus_tag* to extract the gene feature from the genbank file. If the objective is the amplification of an internal region, the *start_position_within_gene* and *end_position_within_gene* options can be used to define the intergenic region to extract. Finally, the promoter and terminator modes, use the cIndex and *locus_tag* to get the gene genomic location, and extract a region of *product_size* length upstream the start codon (promoter mode) or downstream the stop codon (terminator mode).

For the primer design, the *flexibility* option relaxes the region ends so that primers can bind a specified number of nucleotides from the end of the target region. The fixNear or fixFar options indicate whether a primer location should be fixed, allowing only its length to vary. The FixedNear and FixedFar options behave differently across region, gene, promoter, and terminator modes. In region, gene, and promoter modes, FixedNear anchors the primer to the start codon side (right in region mode). In terminator mode, it fixes the primer to the stop codon side. This positioning is relative to the gene’s coding strand (plus or minus). Once the target sequence has been defined, the program produces a restriction enzyme report. This report identifies recognition sites for a list of user-specified REs within the sequence, aiding in the selection of RE for downstream PCR cloning. Additionally, 5’-oligonucleotides –including RE recognition sites– can be configured to be added to the primer design by Primer3.

The primer design performed using Primer3 incorporates configuration parameters such as Tm and product size ranges, the oligonucleotides to add to the primers, the target DNA sequence, and whether the primers should be fixed on either side of the template (Figure 2.A-B). A list of the 5 best primer sets is reported.

Finally, the designed primers are used as a query for BLASTn and searched against the genome as target. BLASTn is run with the task *blastn-short* and word size of 4 to allow for the detection of the primers, including imperfect hits. As a result, two tables are generated for each set of primers, one containing the raw results of blast, and the other reporting primer pairs which could amplify a fragment in a PCR. The first table is filtered so that only the primers for which the BLAST alignment length is at most 3 bases shorter than the primer length are reported. For the second table, the BLAST hits for each forward and reverse primer set are analyzed in pairs to find primers that could produce a PCR product—i.e. primers binding in opposite strands and facing the same direction— of less than 20.000 nucleotides, and considering both perfect and imperfect matches (Figure 2.B).

### Validation of the designed primers

ColabPCR was employed to design primers for the amplification of putative promoter and terminator regions from *Daucus carota* genes, utilizing the NCBI GCA_001625215.2 assembly accession as a template. To ascertain the success rate of the primers generated by ColabPCR, an experimental validation was conducted. The designed primers were synthesized, and a PCR reaction was performed using genomic DNA from *Daucus carota* (Ergo Bioscience SAS proprietary cell lines) as a template. Three batches of primers were designed and evaluated. The first batch consisted of primers generated with the program’s optimal score, without virtual PCR for specificity assessment. For the second batch, a virtual PCR analysis was performed to ensure 1:1 specificity between forward and reverse primers and the absence of mismatches, guaranteeing exclusive binding to the target sequence.

Primers for amplification were ordered (Microsynth AG) and PCR reactions carried out using Q5 DNA Polymerase (New England Biolabs) according to manufacturer’s instructions. Melting temperatures were calculated using the manufacturer’s Tm calculator (https://tmcalculator.neb.com/). In total, 38 PCR reactions were carried out. To determine whether amplifications were successful, the PCR products were analyzed via agarose gel electrophoresis.

A notable enhancement of almost 10% was recorded in the second primer batch, as detailed in Table 1. Furthermore, the second primer design batch was successful in targets that couldn’t be amplified in the first attempt. This significant improvement underscores the effectiveness of a meticulous primer selection strategy. Specifically, prioritizing primer sets that are rigorously screened to be free of unintended secondary binding sites and are structurally optimized for efficient PCR product generation proves to be a highly advantageous methodology.

**Table 1.** Evaluation of the primer designed using ColabPCR.

| Primer Batch | Success Rate | Number of Reactions |
| --- | --- | --- |
| 1 Primers with best score | 62% | 24 |
| 2 Primers with best score<br>Evaluation by Virtual PCR | 71% | 14 |
| 3 Primers with best score<br>Evaluation by Virtual PCR<br>Automated cloning platform | 74% | 83 |

This approach not only minimizes the risk of off-target amplification and primer-dimer formation but also maximizes the specificity and yield of the desired amplicons. Consequently, adopting such a refined primer design and validation process is a crucial step towards consistently increasing the overall success rate of PCR experiments, leading to more reliable and reproducible results in molecular biology applications. Furthermore, the existing Primer design programs do not provide a comprehensive report on potential secondary targets that might result in a PCR product.

These optimized conditions for PCR primers design and the verified increase in amplification success rate allowed us to partner with an automated cloning and assembly platform (Global Institute for Food Security, GIFS, Saskatoon, Canada) to test our design strategy with a larger number of genomic elements. A third primer batch targeting both promoter and terminator regions was therefore designed (a restricted flexibility setting at one of the target ends was used) incorporating restriction enzyme sites compatible with the cloning strategy at the 5’ end of the primers. Selection of the cloning sites was guided by the restriction enzyme report generated by ColabPCR. The amplification rate for this third primer batch was slightly increased, yielding 83 amplicons of the expected size out of 112 attempts (74% success rate, Table 1). These amplicons were sequentially purified, digested, and assembled using restriction endonucleases, resulting in an overall correct assembly and identity-confirmation efficiency of 68% based on whole-plasmid sequencing). For eight specific elements in which amplification failed in the initial attempt, additional primer pairs were designed by modifying the initial settings and/or target region. Of these 8 specific elements, 4 were successfully amplified, being only one a perfect match with the reference genome. This strongly suggests that primer design is not the primary cause of amplification failure and that other factors, such as discrepancies between the chosen carrot variety and the reference genome, may influence the outcome.

## Conclusions

ColabPCR is a Python notebook that offers a streamlined and reproducible workflow for primer design and evaluation in molecular biology. It centralizes several key functionalities within a single interface, utilizing Google Colab’s computational resources to ensure high performance and accessibility without requiring local software installation. While it shares features with other tools—such as using Primer3 for primer design and employing virtual PCR for evaluation—ColabPCR provides distinctive capabilities. These include a clear report of off-target primer binding sites (Figure 2.A-B) and a comprehensive analysis of primer orientation and inter-primer distance to predict PCR product formation, which helps in selecting the most efficient primer set and reducing reaction inefficiency. Its core capabilities extend to simplifying the extraction of the target sequence from the genome to automate the design and evaluation process, primer design, restriction enzyme site verification within the target sequence, incorporating modifications for downstream cloning, performing whole-genome off-target analysis. All these features contribute to a simpler, more accurate, and ultimately more successful primer design process.

## Author contributions

M.J.L: Conceptualization, Methodology, Software, Visualization, Writing-Original draft preparation, Writing-Reviewing and Editing. D.D: Investigation, Validation, Writing-Original draft preparation. G.B: Investigation. J.S: Investigation. J.K-B: Investigation. E.J.P: Conceptualization, Investigation, Formal analysis, Writing-Reviewing and Editing, Project administration, Funding acquisition. All the authors critically reviewed the submitted version of the manuscript.

## Acknowledgments

ColabPCR was developed by M.J.L, D.D, E.J.P, and Ergo Bioscience SAS. M.J.L is affiliated with CONICET (Consejo Nacional de Investigaciones Científicas y Tecnológicas, Argentina) and UNLP (Universidad Nacional de la Plata) as a researcher. This research was carried out at the IBBM (Instituto de Biotecnología y Biología Molecular), an institute jointly managed by CONICET-CCT La Plata and UNLP. E.J.P holds a research position at CONICET, based at the IBBM, and is the CSO and Cofunder of ERGO Biotechnology. D.D is an employee of ERGO. G.B, J.S and J.K-B are employees of NOVABIOMA Biomanufacturing and are affiliated to ACIB GmbH. We acknowledge Benjamin Scott and Megha Bajaj from GIFS (https://gifs.ca/) for their support with the automated cloning platform.

## Data availability statement

The source code for ColabPCR is available on GitHub at https://github.com/maurijlozano/ColabPRC and Zenodo https://doi.org/10.5281/zenodo.18034799 as a Jupyter notebook. The software is distributed under the MIT license.

